# Spatio-temporal dynamics of the proton motive force on single bacterial cells

**DOI:** 10.1101/2023.04.03.535353

**Authors:** Anaïs Biquet-Bisquert, Baptiste Carrio, Nathan Meyer, Thales F.D. Fernandes, Manouk Abkarian, Farida Seduk, Axel Magalon, Ashley L. Nord, Francesco Pedaci

## Abstract

Electrochemical gradients established across biological membranes are fundamental in the bioenergetics of all forms of life. In bacteria, the proton motive force (PMF), the electrochemical potential associated to protons, powers an impressive array of fundamental processes, from ATP production to motility. While far from equilibrium, it has classically been considered homeostatic in time and space. Yet, recent experiments have revealed rich temporal dynamics at the single cell level and functional spatial dynamics at the scale of multicellular communities. Lateral segregation of supramolecular respiratory complexes begs the question of whether spatial heterogeneity of the PMF exists even at the single cell level. By using a light-activated proton pump as a spatially and temporally modulatable source, and the bacterial flagellar motor as a local electro-mechanical gauge, we both perturb and probe the PMF on single cells. Using global perturbations, we resolve temporal dynamics on the ms time scale and observe an asymmetrical capacitive response of the cell. Using localized perturbations, we find that the PMF is rapidly homogenized along the entire cell, faster than proton diffusion can allow. Instead, the electrical response can be explained in terms of electrotonic potential spread, as found in passive neurons and described by cable theory. This implies a global coupling between PMF sources and consumers in the bacterial membrane, excluding a sustained spatial heterogeneity while allowing for fast temporal dynamics.

**Significance:** Storing energy in the form of a proton gradient across a membrane is a fundamental feature of living systems. In mitochondria, spatial compartmentalization separates electrically distinct regions. In bacteria, it is unclear how this energy reservoir, the proton motive force, behaves at the single cell level: can it be heterogeneous in space as in mitochondria? How fast can it change in time? Using a light-driven proton pump and the flagellar motor as a local electro-mechanical gauge, we find that the bacterial proton motive force can change in a few tens of milliseconds, and that it is instantaneously homogenized along the membrane. This electrophysiological response is surprisingly similar to electrotonic voltage spread in passive neurons.

## 1 Introduction

Evolved in all kingdoms of life, electrochemical gradients established out of equilibrium across bioenergetic membranes are a universal intermediate in energy conversion. Protons play a major role in processes such as cellular respiration and photosynthesis, leading to ATP synthesis, wherein proton translocating enzymes generate a concentration gradient across specialized membranes which act as effective capacitors. The resulting stored electrochemical potential energy, the proton motive force (PMF), is composed of a difference in proton concentration and electrical potential across the membrane, as given by Mitchell’s classical chemiosmotic theory [1].

While long believed to be constant in time and space along the membrane, there is growing evidence pointing towards a rich spatio-temporal dynamical behavior of the PMF, though the biologically functional consequences remain poorly understood [2]. In mitochondria, where the PMF drives ATP synthesis, recent evidence points to a local and dynamic PMF heterogeneity, arising from the complex geometry of the inner mitochondrial membrane, supramolecular organization, and microcompartmentation [3–5]. In bacteria, where the PMF is known to regulate ATP synthesis [6], cell division [7], motility [8], antibiotic resistance [9], sporulation [10], and biofilm formation [11], recent observations also suggest a rich dynamical behavior. Rapid membrane depolarizations observed on single *Escherichia coli* cells [12], related to mechanosensing [13] and inter-cellular communication within biofilms [11], are indications of a temporally dynamical PMF. Moreover, it is well established that oxidative phosphorylation complexes dynamically organize into supramolecular micro-domains [14, 15]. Such spatio-temporal organization has been suggested to be beneficial for the stability of the individual complexes, to confine mobile electron carriers, and to limit the production of reactive oxygen species. By bringing proton consumers and producers close together, this opens up the possibility for compartmentalization of the proton cycle, locally altering the PMF. However, the question of whether a spatially heterogeneous distribution of proton sources and sinks can produce a heterogeneous PMF for specific functions in living cells remains open [14–18].

Despite its importance, the spatial and temporal dynamical behavior of the PMF in bacteria remains poorly characterized and understood, especially at the single cell level. The challenge lies in bacteria’s micrometer-scale and their complex membrane composition, which precludes the use of classical techniques such as patch clamp, except at the expense of large perturbations to the cell envelope [19, 20]. Techniques such as Nernstian and pH-sensitive dyes as well as genetically-encoded pH and voltage sensors, have recently enabled quantitative measurements at the single cell level, pushing forward the nascent field of bacterial electrophysiology [12, 21, 22]. While extremely valuable, these methods come with difficulties in transformation or labelling, as well as calibration of the often complex fluorophore response.

In this work, we turn to a biological probe to characterize the spatio-temporal PMF dynamics on single *E. coli* cells. The bacterial flagellar motor (BFM) is the PMF-driven nano-rotary motor located at the base of each flagellum, which drives cell motility in many motile bacteria [21, 23, 24]. We use spatio-temporally structured laser excitation on single bacteria expressing the light-driven proton pump proteorhodopsin (PR) to controllably augment the PMF in space and time. Taking advantage of the linearity between BFM rotational speed and PMF [25–27], and building on previous works [27–32], we use the speed measurement of individual BFMs to resolve local PMF dynamics in the millisecond range on single cells.

Our measurements show that the capacitive response of the cell is asymmetrical and allows temporal PMF variations on the ms time scale, while spatially localized PMF perturbations are effectively homogenized over the entire cell. Such spread of the PMF occurs faster than diffusion can allow, in a process that links the electrical activity of bacteria with the electrotonic response of passive neurons, quantitatively explained by cable theory [33, 34]. This implies that a sudden increase in PMF due to a localised source is experienced synchronously by all sinks on the membrane, prompting a global response to a local perturbation. Such global coupling between sources and sinks, and the consequent fast PMF homogenization, should exclude the presence of a functional heterogeneity of the PMF resulting from membrane compartmentalization.

## 2 Results

Our measurements were based on a modification of the BFM bead assay, typically employed to study the dynamics of the motor [24, 35]. We measured the BFM angular speed via bright field microscopy (with up to a 20 kHz sampling rate [36]) by tracking the recorded holograms of the off-axis rotation of 600 nm-diameter beads attached to flagellar stubs of individual immobilized *E. coli* cells (see Fig. 1a, and Materials and Methods for details). We simultaneously manipulated the PMF of PR-expressing cells via laser excitation (see Materials and Methods and Supplementary Information section 1 for details), producing a local excess of external proton concentration controlled by the the spatio-temporal profile of the laser beam.

**Figure 1:**
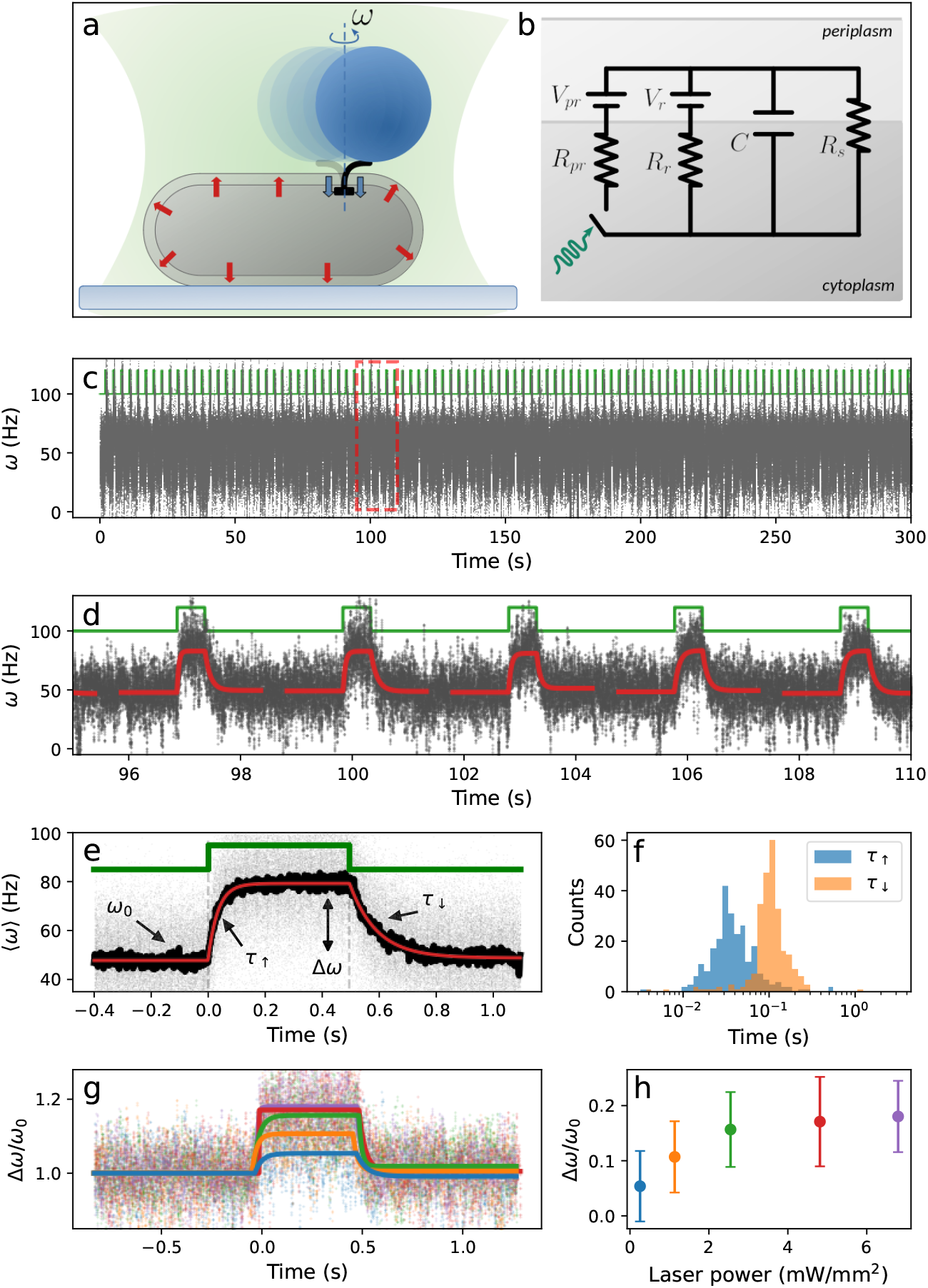
Temporal PMF dynamics on a single *E.coli* cell, probed by BFM rotation and PR excitation. (a) Schematic of the experimental assay. An E. *coli* cell attached to a coverslip and a 600nm bead bound to the a truncated flagellar filament enabled measurements of the BFM speed response to PMF changes triggered by laser-activated PR. Blue inward arrows indicate the PMF consumption by the BFM and red outward ones, the PMF generated by the laser-activated PR. (b) Circuit model describing the cell. PR and respiratory complexes are modeled as voltage sources *Vp_r_, V_r_* with internal resistances *Rp_r_, R_r_*. The membrane has capacitance *C* and PMF sinks are modeled as a resistance *R_s_*. PR laser excitation opens and closes the switch. (c) BFM speed response (grey) induced by a series of on-off rectangular laser pulses (green). (d) A zoom from a region in (c) indicated by red dashed lines. A piecewise function composed of single exponentials (Eq. 6, Materials and Methods) was fit to each speed transition generated by a laser pulse (red). (e) The overlay of 100 laser-synchronised transitions (grey), the mean response (black), and the fit of the mean (red). The motor speeds up upon a laser pulse from its steady-state speed *ω*_0_ to a excited steady-state speed *ω*_1_. *τ*_↑_ and *τ*_↓_ are the characteristic relaxation times. (f) Histogram of the characteristic times, *τ*_↑_ and *τ*_↓_, extracted from the fit of each speed transition shown in (c). (g) The average (dots) and fit (line) of dozens of normalized speed transitions on a single motor, where different colors show different laser powers. (h) The increase in speed as a function of laser power, extracted from (g)

### 2.1 PMF temporal dynamics

We first probed the temporal response of the system by monitoring BFM speed during periodic, spatially homogeneous laser illumination events. Under the assumption that PR is homogeneously distributed in the inner membrane, laser illumination creates a homogeneous excess of external proton concentration, globally increasing the PMF of the cell under study. At each excitation pulse, we observed the motor accelerating from the native steady-state speed *ω*_0_ to a higher speed *ω*_1_, due to the PR-generated PMF increase (Fig. 1c-e). We note that the number of the active stator units in the motor is dynamic, mechanosensitive, and PMF dependent [24, 29, 37, 38]. By tuning the laser pulse duration and the time between pulses, we aimed to avoid an increase in the time-averaged PMF and thus avoid an increase in stator number. We recorded up to thousands of transitions from *ω*_0_ to *ω*_1_ on single motors (see Fig. 1c), where 〈*ω*_0_〉 (and thus stator number) remained constant. Traces which showed obvious stator incorporation or dissociation events were manually excluded; while the case of variable stator stoichiometry is interesting, it is left for future studies.

By overlaying and averaging hundreds to thousands of laser-synchronised speed transitions, we obtained the average PR-induced speed response with high signal-to-noise ratio (Fig. 1e, black line), which strongly evokes the charge and discharge of a capacitor. Following previous works [28, 30, 32, 39], we thus modeled the cell by a single lumped electric circuit (Fig. 1b), where the inner membrane acts as a capacitor separating the the cytoplasm and periplasm (we assume the periplasm and extracellular volume to be in thermodynamic equilibrium due to the high permeability of the outer membrane [40]). Analogous to a voltmeter, the BFM produces a speed proportional to the voltage across the capacitor. Proton sinks in the membrane (e.g. proton channels like ATP synthase) act as resistances in parallel, and are combined in the resistance *R_s_*. Native membrane-embedded respiration complexes, which act as proton sources, are modeled as a single voltage source *V_r_* with internal resistance *R_r_*. Similarly, once excited, PR channels act as a light-controlled proton source, modeled by a light-triggered switch in the circuit with a voltage source *V_pr_* and internal resistance *R_pr_*. The analytical expression for the transmembrane voltage, established across the capacitor and detected by the BFM (solved from the circuit in Fig. 1b), can be written by separating the charging *V*_↑_(*t*) and discharging *V*_↓_(*t*) processes, upon inclusion of the PR arm under laser excitation, as,

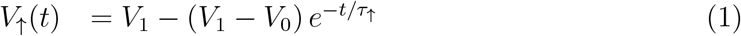

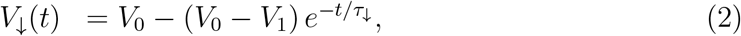

where 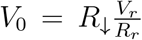 and 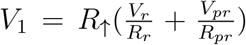 are the native and excited steady-state voltages, respectively, with equivalent resistances 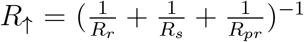 and 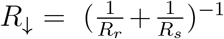, and where the characteristic relaxation times can be written as *τ*_↑_ = *R*_↓_*C* and *τ*_↑_ = *R*_↑_*C*, for the charging (↑) and discharging process (↓), respectively. We note that the solution predicts two distinct relaxation times for the charge and discharge, and that *τ*_↑_ > *τ*_↓_. This can be understood by the fact that the arm of the circuit associated to PR, present only during the charging due to laser illumination, brings the extra resistance *R_pr_*, which makes the equivalent resistance *R*_↑_ < *R*_↓_.

The experimental mean response of the motor to the pulsed PMF excess was fit by a continuous piece-wise function built with *V*_↑_(*t*) and *V*_↓_(*t*) as shown by the red line in Fig. 1e (each synchronized with their relative laser pulse edges, such that at the laser-off time the two functions are equal, see Methods). In agreement with the prediction from the electric circuit model, we find that *τ*_↑_ > *τ*_↓_, as shown by a fit of the the mean transition in Fig. 1e (black line, *τ*_↑_ = 92 ± 0.5 ms and *τ*_↓_ = 36 ± 0.5 ms, see SI section 3 for uncertainty estimation). This feature was robustly observed in all the measured motors (total of 12 motors, see global statistics in SI section 4), and is in direct agreement with the prediction of the electric circuit model. Fits of each individual transition, although noisier, also show this trend (Fig. 1f). In Fig. 1g-h we show that the speed change Δ*ω* increases with laser power, up until a saturation, as seen previously [28]. This could either be explained by the fact that *R_pr_* decreases to a finite value with light intensity, reflecting the limiting rate of PR pumping [28, 30], or by an intrinsic saturation in the motor. Analogously, oxygen depletion, which limits proton pumping of respiratory complexes (see Methods), can be modeled by an increasing *R_r_*. The model shows that, in this case, the difference *V*_1_ – *V*_0_, proportional to Δ*ω*, increases in line with our observations.

### 2.2 PMF spatial dynamics

We then asked whether a spatial heterogeneity of the PMF can exist and be detected by our measurements. To maximise the possible effect of an inhomogeneous PMF, we used filamentous and PR-expressing *E. coli* cells with lengths up to 30 μm (see Materials and Methods for details), which we periodically excited at only one pole, using a 8 μm diameter laser spot. We chose filamentous cells where two functional motors were labeled by beads, giving two PMF reporters at different distances from the proton source (up to a max of 20 μm), which was spatially and temporally defined by the intersection of the laser beam with the cell (Fig. 2a,b). As a control, we recorded the two motor speeds during localized illumination of each cell pole, such that each motor played the role of both the distal and proximal sink. The measurements (Fig. 2a-f) show that both motors display the same charging-discharging behavior described above, with a temporal profile that is strikingly similar, within error, independent of their distance from the source. Two features, in particular, can be highlighted: i) the response of both motors starts at the laser-on edge with no resolved delay (within our time resolution of 2 ms, see SI section 2), and ii), once normalized to the steady state value *ω*_0_, the speed increase is the same for both motors (Fig. 2c,f). Both features were observed in all measurements (seven filamentous cells), as summarized in Fig. 2h. We conclude that an excess of PMF generated locally can propagate for tens of micrometers in less than a few milliseconds with no measurable loss, such that all sinks within these distances are affected identically.

**Figure 2:**
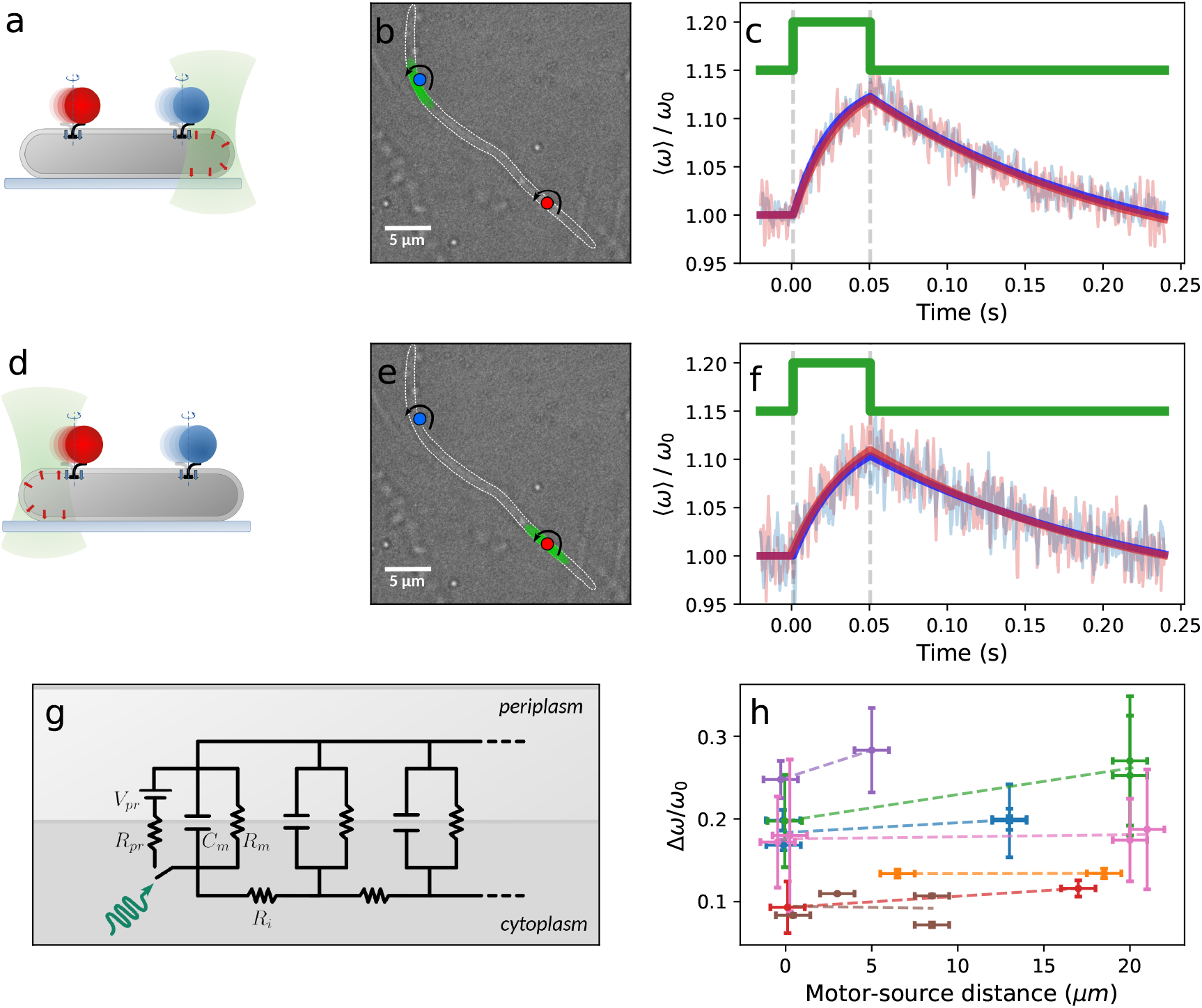
Spatial PMF dynamics. (a) Local illumination of a filamentous *E. coli* cell, where activating PR triggers a local PMF increase. The speed response of two motors located close (blue) and far (red) from the source was measured. (b) Microscope image of the assay depicted in (a), where the green spot indicates the portion of the cell illuminated by the laser and colored circles indicate the rotating beads attached to truncated filaments (c) Average of 1493 (red) and 1793 (blue) laser-synchronised speed transitions (faint lines) and the fits (dark lines) of the two motors shown in (b) located near (blue) and far (red) from the laser (green). (d-f) Same as (a-c) with the local illumination at the opposite motor. (f) Average of 751 (blue) and 690 (red) transitions. (g) *E. coli* circuit model based on cable theory and consisting of sub-circuits in series, each modeling a small section of the cell. A non-illuminated module is represented by the membrane capacitance (*C*) and sinks (*R_s_*) whereas the illuminated module includes the PR arm as for the circuit in figure 1a. (h) Relative increase in speed as a function of the distance between the source and the motor, where measurements on motors of the same cell are indicated by points of the same color, for which a linear fit (dotted line) is shown to guide the eye.

We then asked whether the PMF generated on one cell can be transmitted to cells in close proximity. We selected pairs of non-filamentous cells located less than 8 μm from each other, both with a bead-labeled functioning motor. Periodically illuminating only one cell at a time, we observed that only the motor belonging to the illuminated cell increased its speed following the laser (Fig. 3a-d). Similarly, we sequentially illuminated motors on two overlapping filamentous cells, and we observed that only the motor of the illuminated cell showed PR-induced acceleration (Fig. 3eh), irrespective of the laser position along the cell. These observations indicate that the PMF excess produced across one membrane cannot power a motor on a separate cell, even if the two cells are in close proximity or in contact.

**Figure 3:**
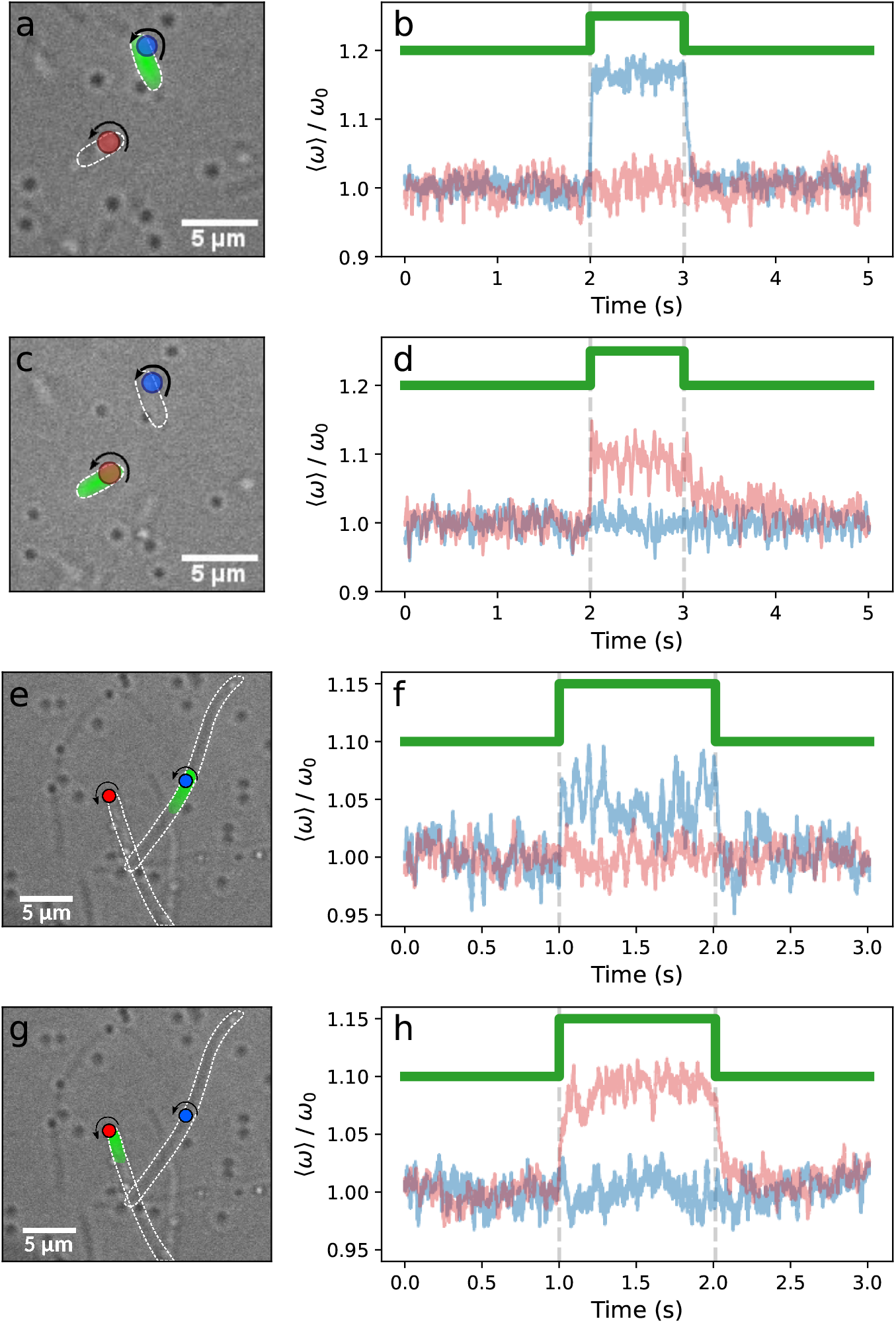
The effect of PR excitation on neighbouring and overlapping cells. (a) Two bacteria, each with a labeled motor, are separated by ~ 3 μm; the cell labeled by the blue bead is illuminated with the laser (depicted in green). (b) Normalized average speed response of hundreds of transitions of the two motors, labeled by the same colors. (c-d) Same as (a-b) but the laser excites the cell labeled by the red bead. (e-f) Image and speed traces for two filamentous overlapping cells, each with a labeled motor. The cell labeled by the blue bead is illuminated by the laser. (g-h) The same as (e-f), but the cell labeled by the red bead is illuminated. The colors of the traces correspond to the color of the bead.

## 3 Discussion

In the first part of this study, we probed the temporal response of the system via homogeneous PR excitation, showing that PR can perturb the PMF, and that the BFM can report upon PMF dynamics, both at the millisecond timescale. We observed relaxation times (*τ*_↑_, *τ*_↓_) on the order of tens of milliseconds, one to three orders of magnitude lower than previous measurements [28, 30, 31]. As it is now known that the number of bound and active stators is proportional to the PMF [21], it is likely that the previously measured timescales were dominated by dynamic stator exchange. By carefully avoiding the effect of stator dynamics, the shorter time scale we resolve reports upon the cellular electrical response. This time scale is unaffected by proton pumping by PR (with the laser off, *τ*_↑_ is not influenced by PR), nor is it severely limited by the elasticity of the flagellar hook (see Material and Methods). The electric circuit model of Fig. 1b predicts an asymmetry in the two characteristic times of the system, which we systematically observe in our experiments (i.e. *τ*_↑_ ≃ 100 ms, *τ*_↓_ ≃ 30 ms, see Fig. 1f). By using the measured values of *τ*_↑_, *τ*_↓_, *ω*_0_ and *ω*_1_, together with an estimate of the membrane capacitance (*C* ≃ 10^-14^*F*) and of the respiration source (*V_r_* ≃ 360 mV) [28, 41], we use the analytical results obtained from the theoretical solution of the circuit to determine the values of all the cellular electrical parameters (see SI sec.5): *V_pr_* ≃ 62 mV, *R_pr_* ≃ 10^12^ Ω, *V_r_* ≃ 360 mV [28], *R_r_* ≃ 9 · 10^13^ Ω, *R_s_* ≃ 10^13^ Ω, *C* ≃ 10^-14^*F* [28, 41], *V*_0_ ≃ 37 mV, *V*_1_ ≃ 60 mV.

In the second part of this work, by controlling the spatial position of the PMF source, we observed that the response of the BFM is independent of its distance from the proton source, up to ~ 20*μm*. It is worth noting that among ions, protons behave peculiarly [42], and the classical model of a “delocalized” PMF (where protons immediately equilibrate into the bulk solution) has been challenged by numerous studies of proton dynamics along membranes. Despite variations in quantitative results and ongoing discussions about the underlying mechanisms, many observations indicate that protons translocated across membranes can remain confined and in close proximity to the membrane [5, 14, 43–50]. Efficient proton diffusion can then occur in two dimensions with a high diffusion coefficient, comparable to bulk (where protons have the highest diffusion coefficient among ions, *D_H_* = 9 · 10^-5^ cm^2^/s, due to the Grotthuss mechanism [51]). While such a “localized” model of PMF provides a coherent picture of energy transduction, evidence comes mainly from controlled *in-vitro* systems; measurements in vivo, and particularly in bacteria, are lacking. Proton diffusion measurements on synthetic lipid bilayers show a substantial deviation from classical diffusion at short time scales (< 100 ms) [48, 52]. Our measurements could be seen as in line with this “anomaly” at the millisecond scale in living bacteria. As we observe that the transfer of PMF does not occur across neighboring or intersecting cells, our results could be explained by a surface-localized PMF transmission, though the need of a more direct proof remains.

It is interesting to note that, despite their different physical nature, proton concentration *ρ*(*x, t*) and transmembrane voltage *V*(*x, t*) (functions of space *x* and time *t*), both contributing to the PMF, obey the same diffusion equation, which can be generally written in one dimension as

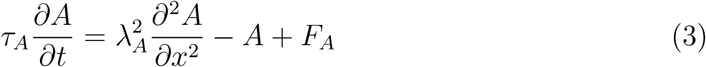

where *A* = *ρ* or *V*, *τ_A_* and *λ_A_* are the characteristic time and length, and *F_A_*(*x, t*) is the source term. Taking *A* = *ρ*, Eq. 3 describes concentration diffusion in the presence of losses from sinks (with first order rate *k_s_*) and gains from sources (with rate *k_pr_*), where the parameters can be expressed as *τ_ρ_* = 1/*k_s_*, 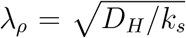, and *F_ρ_* = *k_pr_/k_s_* [52]. On the other hand, taking *A* = *V*, Eq. .3 successfully models the transmembrane voltage in the different context of cable theory, developed for the electro-physiology of unmyelinated passive neuronal axons (with voltage below the action potential threshold) [33, 34, 53]. Here the cylindrical cell is described by an equivalent electric circuit made by serially stacked resistive-capacitive (RC) sub-circuits, each modeling a small section of the cell (Fig. 2g). Solving the circuit, the theory provides the deviation from steady state of the transmembrane voltage *V* [33]. The parameters are here related to the electric components of the circuit as *τ_V_* = *R_m_C_m_*, and 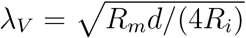, where *R_m_* (units of Ω*m*^2^) is the membrane resistance across a unit area, *R_i_* (Ω*m*) the intracellular volume resistivity, *C_m_* (*F/m*^2^) the capacitance per unit area, d the cell diameter, and *F_V_*(*x, t*) is the source, resulting from an injected current (which, on axons, can be localized by the use of micropipettes).

An analytical solution of Eq. 3 for a spatio-temporally localized source can be found both in the context of diffusion (heat transfer) [54] and in cable theory [34] (see SI sections 6, 7). As shown in Fig. 4, during the charging and as a function of space, the solution peaks at the source location and exponentially decays with characteristic length *λ* (dropping the index for simplicity, see Fig. 4a). During the discharge, the peak is smoothed before the solution reaches zero everywhere at long times. During charging and as a function of time, the solution saturates to a plateau everywhere within a few characteristic times *τ* (Fig. 4b). As the distance from the source increases, both a delay in the rising edge and a decrease in the plateau occur. During discharge, the solution shows an exponential decay, with a delay in the falling edge which increases with distance to the source. We note that the solution of classical cable theory displays symmetric characteristic times during charge and discharge (*τ*_↑_ = *τ*_↓_) due to the fact that the source *F*(*x, t*) is modeled as a current source. We find that by instead modeling the source as a battery *V_pr_* with internal resistance *R_pr_* (Fig. 2g), or by introducing a saturation mechanism in the pumping rate (*k_pr_* = *k_pr_*(*ρ*)) in 2D diffusion simulations, the observed asymmetry is reproduced (see SI sec. 7). This general solution is instrumental to discriminate the relative roles of concentration and voltage in our measurements, as discussed below.

**Figure 4:**
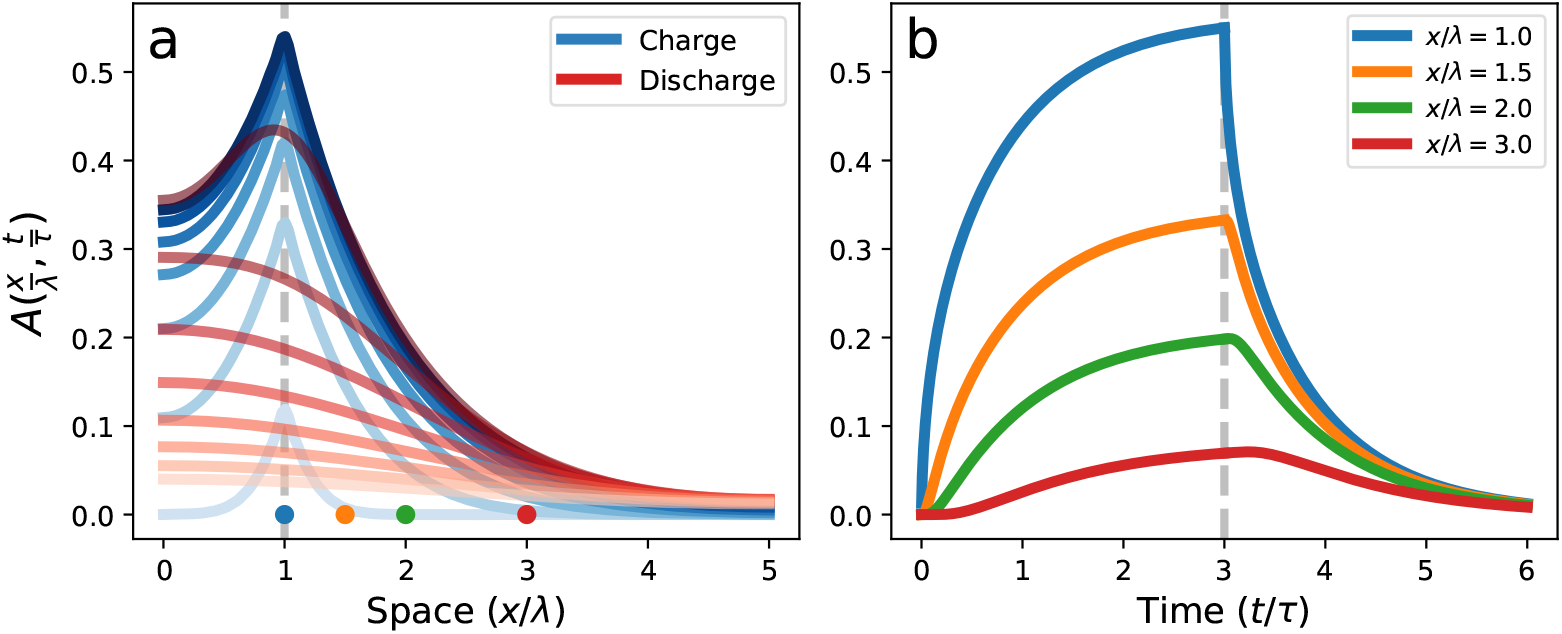
Analytical solution *A* of the cable equation (Eq. 3), function of the normalized variables (*x/λ, t/τ*). The source is a delta-function centered in *x/λ* = 1, active between *t* = 0 and *t* = 3*τ*, and the cell has a length of 5*λ*. a) Behavior of the solution in space, where light to dark blue lines indicate different times during charging (*t* = [0, 3*τ*]), and dark to light red lines indicate different times during discharging (t = [3*τ*, 6*τ*]). b) Temporal profiles of *A*(*x, t*) at positions *x/λ* indicated by the colored dots in a).

Concentration diffusion alone cannot explain our data. In fact, if we assume that the measured characteristic time scale *τ_m_* = 〈(*t*_↑_, *t*_↓_)〉 = (30 – 100) ms is due to concentration diffusion (i.e. *τ_m_* = *τ_ρ_* = 1/*k_s_*), considering the largest diffusion coefficient for protons in bulk (*D_H_*), we estimate the diffusion characteristic length as

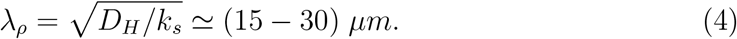

This would set the maximum distance between two experimentally probed motors (~ 20 *μ*m) at about one characteristic length. First, at such a distance, as shown from the analytical solution (comparing the blue and green curves of Fig. 4), we expect a delay in the response of the distal motor on the order of ~ 0.1 *τ_ρ_*, i.e. (3 – 10) ms, which we do not observe within our ms experimental time resolution (see SI section 3). More evidently, we expect a clearly resolvable difference between the plateaus reached during charging between the motor located at the source and the distal motor (30 – 50%, see the blue, orange, and green lines in Fig. 4b). Instead, we do not observe this difference, and the two motors behave identically. We thus conclude that proton diffusion alone is not sufficient to explain the rapid and lossless transmission of PMF.

Instead, if we hypothesize that the measured characteristic time scale *τ_m_* is due to voltage dynamics (i.e. *τ_m_* = *τ_V_* = *R_m_C_m_*), from the values *C_m_* ≃ 1*μF/cm^2^* [28, 41], *R_i_* ≃ 100 Ω *cm* [33, 41], and *d* =1 *μm*, we can estimate *R_m_* ≃ (1 – 10) Ω*m*^2^ and the voltage characteristic length

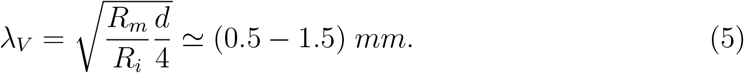

In this scenario, and compatible with our data, the experimental maximum distance between a motor and the PR-induced source (~ 20 *μ*m) is a small fraction of *λ_V_*. Therefore, as the source creates a local proton excess, the transmembrane voltage *V*(*x, t*) takes a few tens of ms (*τ*_↑_) to reach the new steady state, which, although exponentially decaying in space, is nearly constant over the length of cell, evolving in time identically everywhere along the cell. Millimeter-scale characteristic lengths of the transmembrane voltage are also found in passive neuronal axons [33, 34, 53, 55] and confirm the pioneering observations of long distance PMF transmission in mm-scale multicellular filamentous cyanobacteria and mitochondrial networks, performed in the 1980’s by Skulachev and colleagues [56, 57]. The possibility for cells and tissues to develop currents through and around themselves, via asymmetrical distributions of sources and sinks, has also been recognized [58]. At the micro scale, beyond the classical electric circuit description of cable theory, we note that the fact that proton diffusion is not fast enough to explain our observations calls for other mechanisms leading to enhanced proton transport. A complete theoretical understanding of the interplay between charge distribution and transmembrane voltage in the small volume of a bacterium will likely benefit from an electro-diffusion treatment via the Poisson-Nernst-Planck equations, as recognized for mitochondria and sub-micron neuronal compartments [59, 60].

In conclusion, we have found that the bacterial PMF can exhibit temporal dynamics with characteristic times in the tens of milliseconds, depending on the conductance of the ion channels involved. Spatially, our measurements imply that sources and sinks are globally coupled, effectively experiencing the same value of PMF independently of their location in the membrane. This can be explained by the millimeter-scale characteristic length of the transmembrane electrotonic voltage spread from a source over the micron-scale bacterial cell. A spatially homogeneous PMF implies that sinks do not compete for protons and that their consumption rate is independent of their position in the membrane, refuting the hypothesis of local proton circuits in bacteria. For peritrichious bacteria, this feature may in fact be crucial, as theory suggests that in order to form a coordinated flagellar bundle, each of the multiple membrane-bound motors must have nearly equal rotation rates [61]. With similar characteristic length and time scales, despite their vast difference in size, the electrophysiological properties of bacteria and neurons are intriguingly similar.

## 4 Materials and Methods

### Bacteria strain and culture

We used *E. coli* strain MT03, a derivative of RP437 where *cheY* is deleted, producing counter-clockwise only BFM rotation, and *fliC* is replaced with a mutant *fliC^st^*, producing “sticky” filaments and enabling the attachment of polystyrene beads. This strain was transformed with the *pBAD24-PR* plasmid (amp^R^, arabinose induced) containing SAR86 δ-proteorhodopsin. Cells were innoculated from a single colony on a plate into Luria Broth (LB; 10 g L^-1^ bacto-tryptone, 5g L^-1^ yeast extract, 10 g L^-1^ NaCl) containing 100 μg mL^-1^ ampicillin, and grown aerobically overnight at 35 °C, shaking at 200rpm. The overnight culture was diluted to *OD*_600_ of 0.05 with LB, PR expression was induced with 0.01% arabinose, and 10 μM ethanolic all-trans-retinal was added as the necesary cofactor. Cells were grown at at 35 °C and harvested during mid-log growth phase, at an *OD*_600_ of 0.6 – 0.8. Filamentous cells were prepared by adding cephalexin (60 μg mL^-1^) at early-log phase and grown for another 3 h.

### Sample preparation

Flagella were mechanically sheared by passing 1mL of cell culture back and forth between two syringes connected by two 21 gauge needles and a thin tube [26]. Cells were then centrifuged at 3000 rpm for 2 min. After discarding the supernatant, the cells were washed and resuspended in motility buffer (MB, 10mM potassium phosphate, 0.1mM EDTA, 10mM lactic acid, pH 7.0). Custom-made tunnel slides were made of two coverslips separated by a piece of parafilm with a ‘tunnel’ cut from its interior. We flushed 100 μL of poly-L-lysine (Sigma-Aldrich P4707) into the tunnel slide, incubated for 2 min, the flushed through 200 μL of MB. The cell suspension (200 μL) was then flushed through the tunnel slide, and cells were allowed to settle on the coverslip for 10 min. Unattached cells were washed out with MB. Finally, 600nm polysterene beads (Sigma-Aldrich LB6, 300 μL of a 1/300 dilution into MB) were flushed through. Beads were allowed to spontaneously bind to the “sticky” truncated filaments, and after about 10 min unattached beads were washed out with MB.

The motor response to the periodic PR excitation (i.e. Δ*ω* = *ω*_1_–*ω*_0_) was observed to increase, while *ω*_0_ decreased, with the time spent by the cells in the confined volume of the flow slide (data not shown). This is in line with previous observations [28, 30], and may be related to a decreased activity of respiratory complexes with a decrease in available oxygen. In support of this, we also observed an increase in Δ*ω* in the presence of low concentrations of NaN_3_, a bacterial respiration inhibitor (data not shown). It has also previously been shown that PR-induced photocurrent increases as the absolute value of the transmembrane voltage decreases [62]. In order to improve the signal-to-noise of the PR excitation induced transitions, we worked in conditions where oxygen was partially depleted: cells were left in the sealed tunnel slide for about 1 – 2 hours (depending on the bacterial concentration in the sample) before starting the acquisitions. Experiments were performed in MB at 22 °C.

### Microscopy

The custom-built microscope used for all experiments is shown in SI section 1 and described in [36]. The sample was continuously illuminated with a 660nm LED (Thorlabs, M660L3) and imaged with a 100×, 1.45 NA, objective (Nikon) onto a CMOS camera (Optronics CL600×2/M) at 5 – 20 kHz, with a pixel size of 88 nm. PR was excited by a 552 nm laser focused onto the back focal plane of the objective after being spatially limited by an iris. The laser diameter at the sample plane, measured by the fluorescence of a thin layer of Rose Bengal (Sigma-Aldrich, 198250-5G), was ~ 8 μm (see SI section 2 for details), and the intensity at the exit of the objective was 108mW mm^-2^ for PMF temporal dynamics experiments and 6mW mm^-2^ for PMF spatial dynamics experiments. Periodic PR excitation was achieved via an acoustooptic tunable filter (AOTFnC-400.650-TN, AA Opto-electronic) to produce a periodic train of on-off rectangular laser pulses (in the temporal dynamics experiments, the cycle lasted 2.5 s with the laser on for 0.5 s, whereas in the spatial dynamics experiments the cycle lasted 250 ms with the laser on for 50 ms). The position of the laser with respect to the bacteria and rotating beads was determined before each measurement by imaging the sample onto an Electron Multiplying Charge Coupled Device (EM-CCD, iXon Ultra 897, Andor, 109nm per pixel) via a removable mirror and using the autofluorescence of the cell as a proxy for the 552 nm illumination. A small percentage of the modulated PR excitation laser was picked out and imaged onto the corner of the CMOS camera via an optical fiber, in order to automatically synchronize the sampling of laser intensity and motor speed. We note that the 660nm illumination light used for bead tracking is within the tail of the PR excitation spectrum, and it increases the BFM speed by ~ 4%. As it remains illuminated for the entirety of our measurements, its effect is negligible.

### Data analysis

All data analysis was performed using custom Labview and Python scripts. The *x*(*t*), *y*(*t*) positions of the rotating bead were determined by using a cross-correlation analysis of the bead image with a numerically generated kernel pattern [63]. The drift of the circular trajectory was corrected by subtracting a spline interpolation of *x* and *y* from their respective raw values. The elliptical trajectories of the beads, assumed to be the projection of a tilted circle, were transformed into circles by stretching the minor axis of the ellipse. The bead angle was calculated as *θ* = tan^-1^(*y/x*), and the angular velocity as *ω* = *dθ/dt*. The speed trace was low-pass filtered by the Savitzky–Golay algorithm (5^th^ order, 2-8 ms window). Motor speed transitions around a laser pulse were fit using a differential evolution fit algorithm with the following piece-wise function:

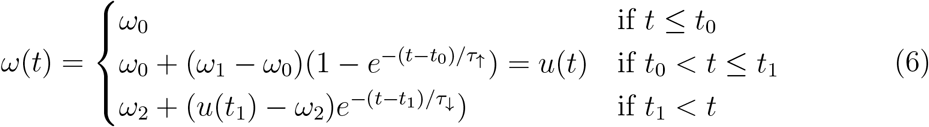

where *t*_0_ (*t*_1_) is the time at which the laser is switched on (off), *τ*_↑_ and *τ*_↓_ are the characteristic times, and *ω*_0_, *ω*_1_, *ω*_2_ are respectively the steady-state speeds before switching on the laser, after switching on, and after switching off. The free parameters of the fit were (*τ*_↑_, *τ*_↓_, *t*_0_, *t*_1_, *ω*_0_, *ω*_1_, *ω*_2_). The distance between the region of the cell illuminated by the laser and the motor was measured using the laser excited auto-fluorescence of the cell and the center of the bead trajectory (see SI section 2 for details). We note that BFM bead assay measurements are intrinsically low-pass filtered by the flagellar hook, the 60 nm extracellular polymer located at the base of the flagellum, which acts as a torsional spring between the motor and the observed load. Given its torsional stiffness [64], the corresponding relaxation time for a 600nm bead tethered to the hook is about 2 ms, which is smaller than the measured characteristic times (*τ*_↑_, *τ*_↓_), and therefore should not severely affect the results.

## Acknowledgments

We thank Emilie Gachon and Florian Morati for preliminary experiments that led to this work. We thank Martin Rieu, Richard Berry, William Hoffman, Nils-Ole Walliser, John Palmeri, Jérôme Dorignac, Fred Geniet, and Andrea Parmeggiani for fruitful discussions. AB-B, AM, FS, ALN, and FP were supported by the ANR Flag-Motor project grant ANR-18-CE30-0008 of the ANR, the French Agence Nationale de la Recherche. TFDF and FP were supported by the ANR HighResPMF project grant ANR-20-CE42-0005-01. ALN was further supported by the ANR PHYBABIFO project grant ANR-22-CE30-0034. The CBS is a member of the France-BioImaging (FBI) and the French Infrastructure for Integrated Structural Biology (FRISBI), two national infrastructures supported by the French National Research Agency (ANR-10-INBS-04-01 and ANR-10-INBS-05, respectively).

## Supplementary Information

### 1 Microscope setup

**Figure 1:**
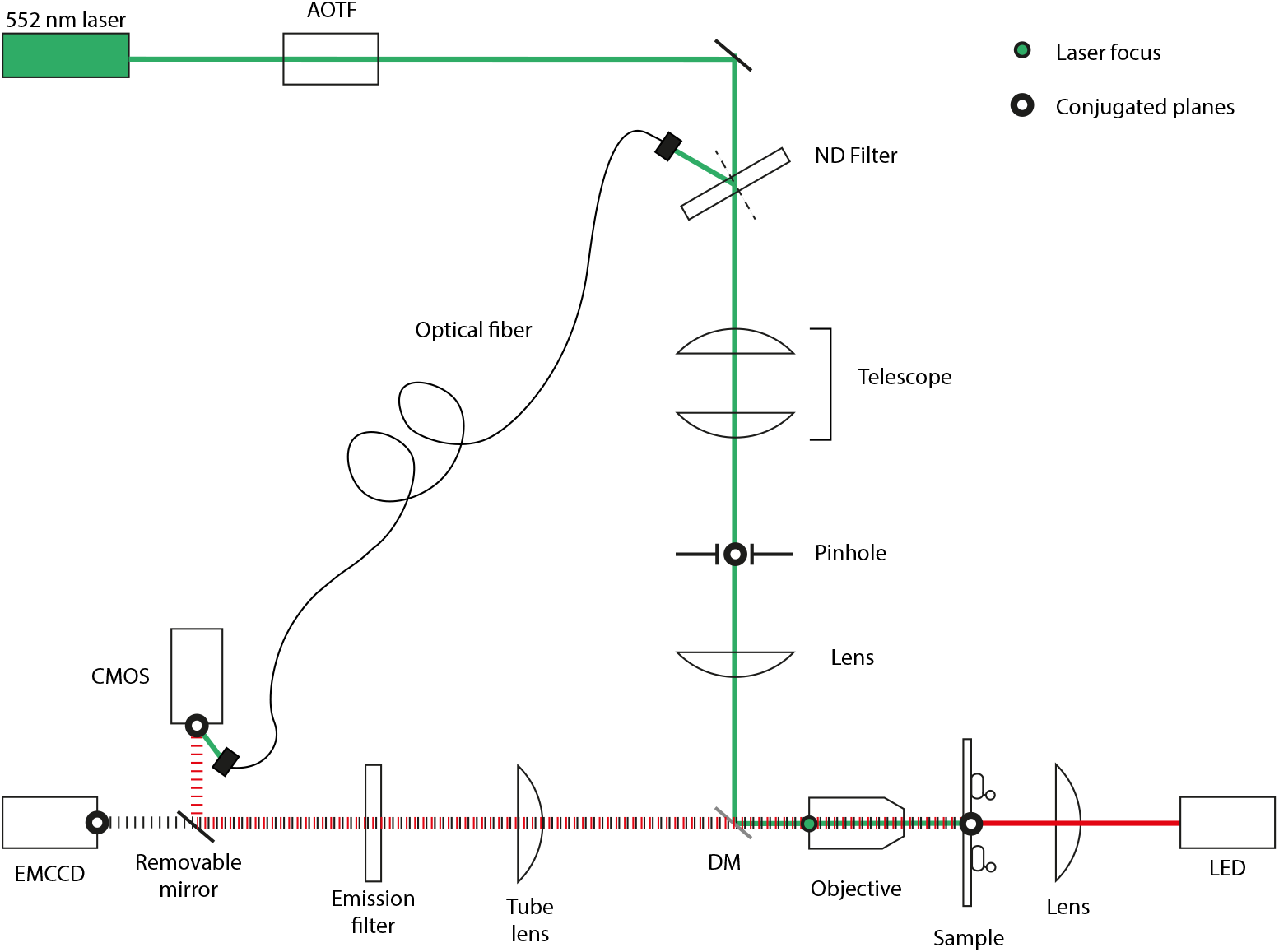
Schematic of the optical setup consisting of two illumination paths: brightfield and epifluorescence. In the first path (red) the sample was illuminated with a 660nm LED and imaged with a 100× (1.45 NA) oil-immersion objective (Nikon) onto a fast CMOS camera (Optronics CL600×2/M). In the epifluorescent path (green), a 552nm laser controlled by an Acousto-Optic Tunable Filter (AOTFnC-400.650-TN, AA Opto-electronic) is used to excite PR. The beam intensity was reduced via an ND filter and its diameter was controlled by lenses and a pinhole. The auto-fluorescence of the cells was imaged via a removable mirror onto a cooled Electron Multiplying Charge Coupled Device (EMCCD, iXon Ultra 897, Andor). Using an optical fiber collecting the reflected light from the ND filter, the laser intensity was imaged onto a corner of the CMOS, providing the state of the laser synchronously with the acquisition of the rotating bead.

### 2 Laser spot size measurement

SI Fig. 2a shows the position of the laser next to a cell at a distance of 4 μm, measured from the center of the laser spot to a rotating motor represented in red. At this distance, we observed no effect of the laser on the motor speed (SI Fig. 2c, red). On the contrary, the speed of the same motor, indicated in blue in SI Fig. 2b-c, follows the switching on and off of the laser when the center of the laser is placed on the cell, again at a distance of 4 μm from the motor. This indicates that the effective laser spot diameter is ~ 8 μm. The laser spot size was also measured using a thin layer of fluorescent dye solution in water. The sample was prepared by placing a drop of 0.3 g L^-1^ Rose Bengal (Sigma-Aldrich, 198250-5G) dye solution, previously sonicated for 30 min, between two cleaned coverslips and letting the solution sit for 10 min. Then, the sample was illuminated with the laser (at a power of 6 mW mm^-2^, the intensity used in the PMF spatial dynamics measurements) and imaged onto the EMCCD camera (SI Fig. 2d,e,f).

**Figure 2:**
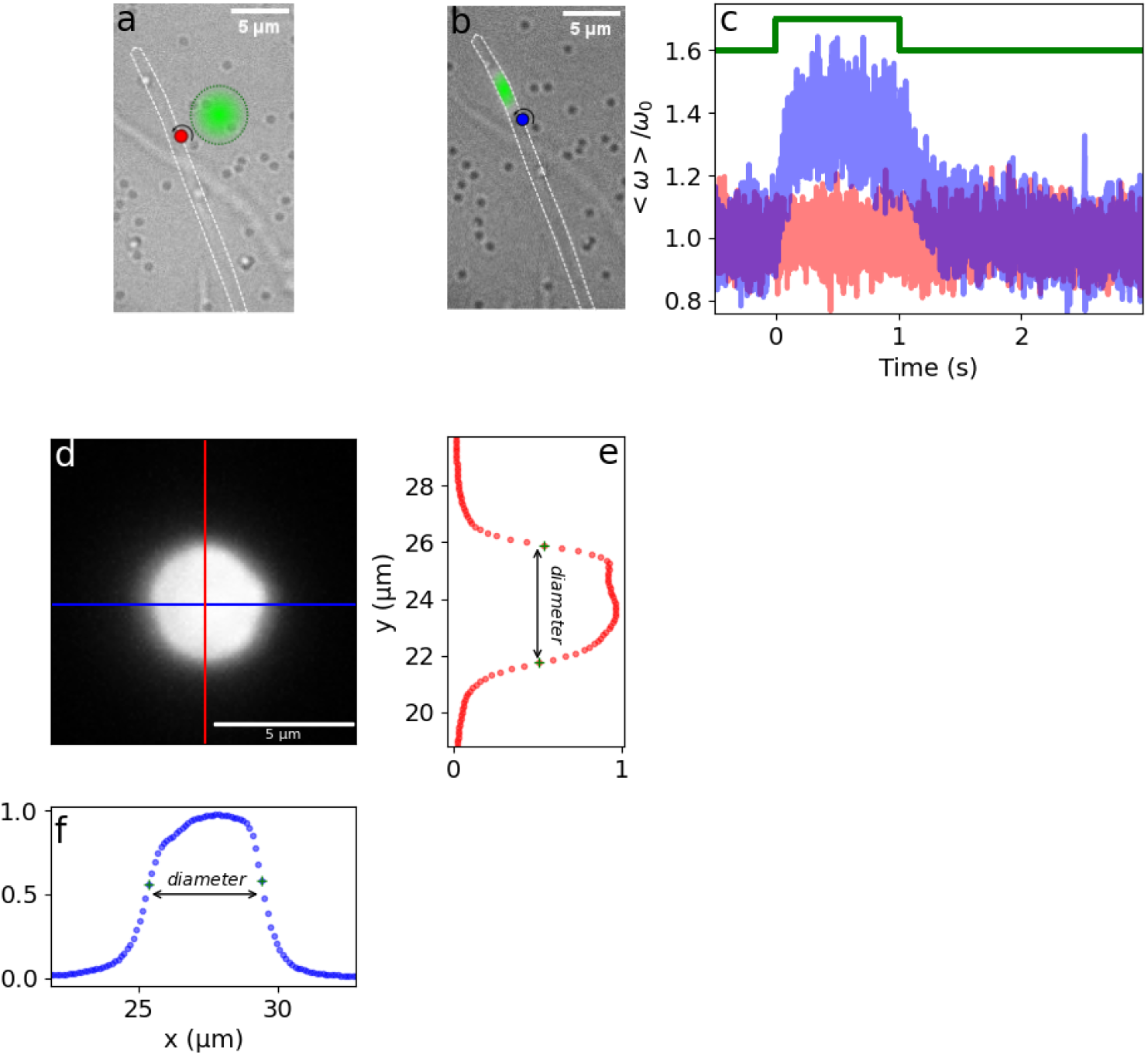
Controls. (a) image of a filamentous *E. coli* cell (outlined in white) with a functioning motor labeled by a bead (in red). The laser (in green) is placed on the coverslip at a distance of 4 μm from the motor. (b) The laser spot is moved on the cell, 4 μm from the bead. (c) Normalized average speed response for 58 transitions of the motor from configuration (a) in red and configuration (b) in blue. The speed response was low-pass filtered by the Savitzky–Golay algorithm (5^th^ order, ~ *6ms* window). (d) Image of a thin layer of fluorescent Rose Bengal solution illuminated with the laser using a CCD camera. (e-f) Corresponding vertical and horizontal intensity profiles.

### 3 Error estimation

**Figure 3:**
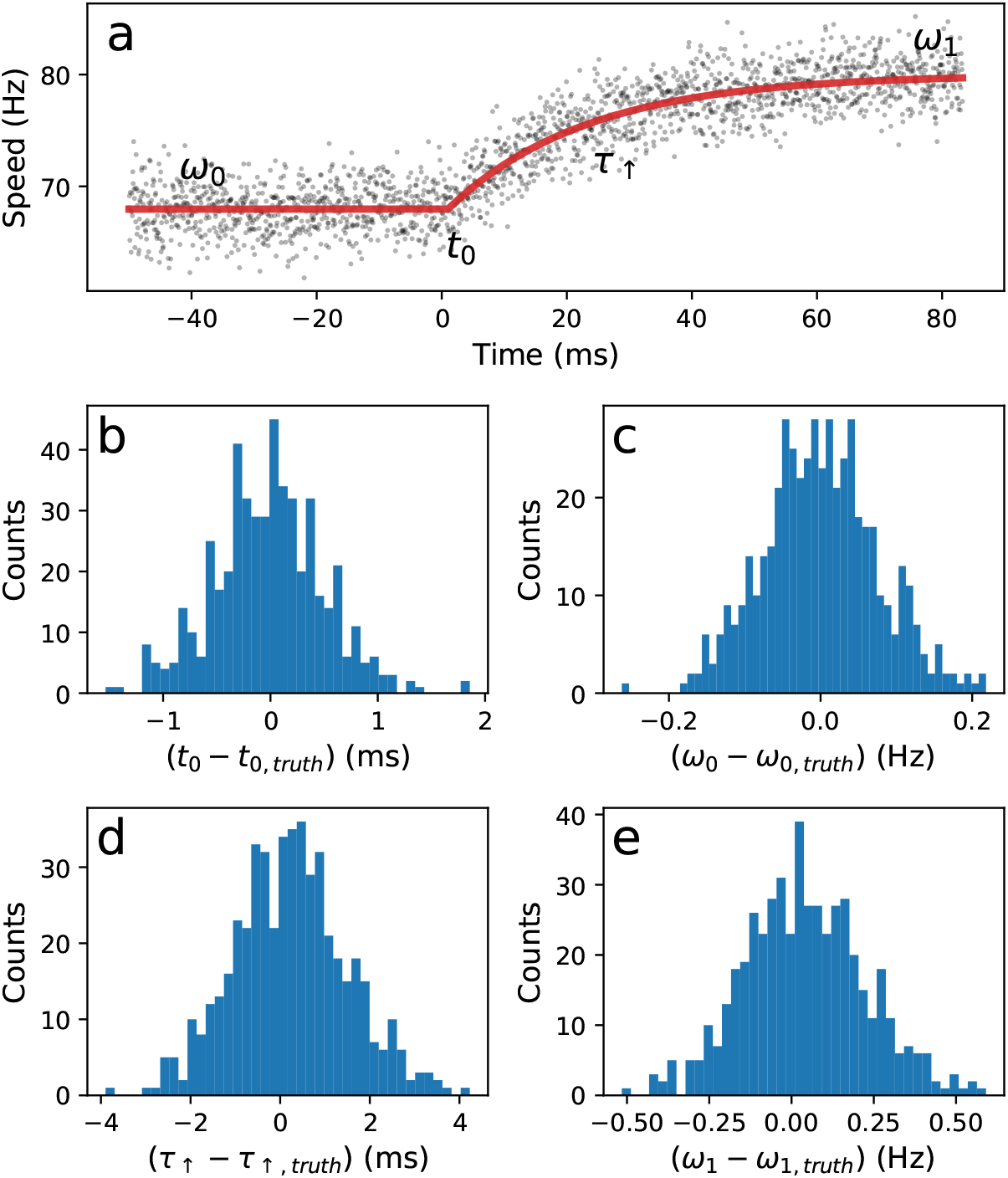
Resolution estimation.(a) A synthetic trace (dark points) is created mimicking the experimental charging response at the laser-on edge, using a constant value followed by an exponential charging function (red line, see Eq.6 of the main text), to which a gaussian noise is added, with standard deviation equal to the one of the experimental trace. The fitting algorithm is challenged against 500 different instances of the noise. The distance distributions between the fitted parameters (*t*_0_, *ω*_0_, *τ*_↑_, *ω*_1_) and the corresponding true ones, are indicated in panels (b),(c),(d), and (e), respectively.

### 4 Global statistics

**Figure 4:**
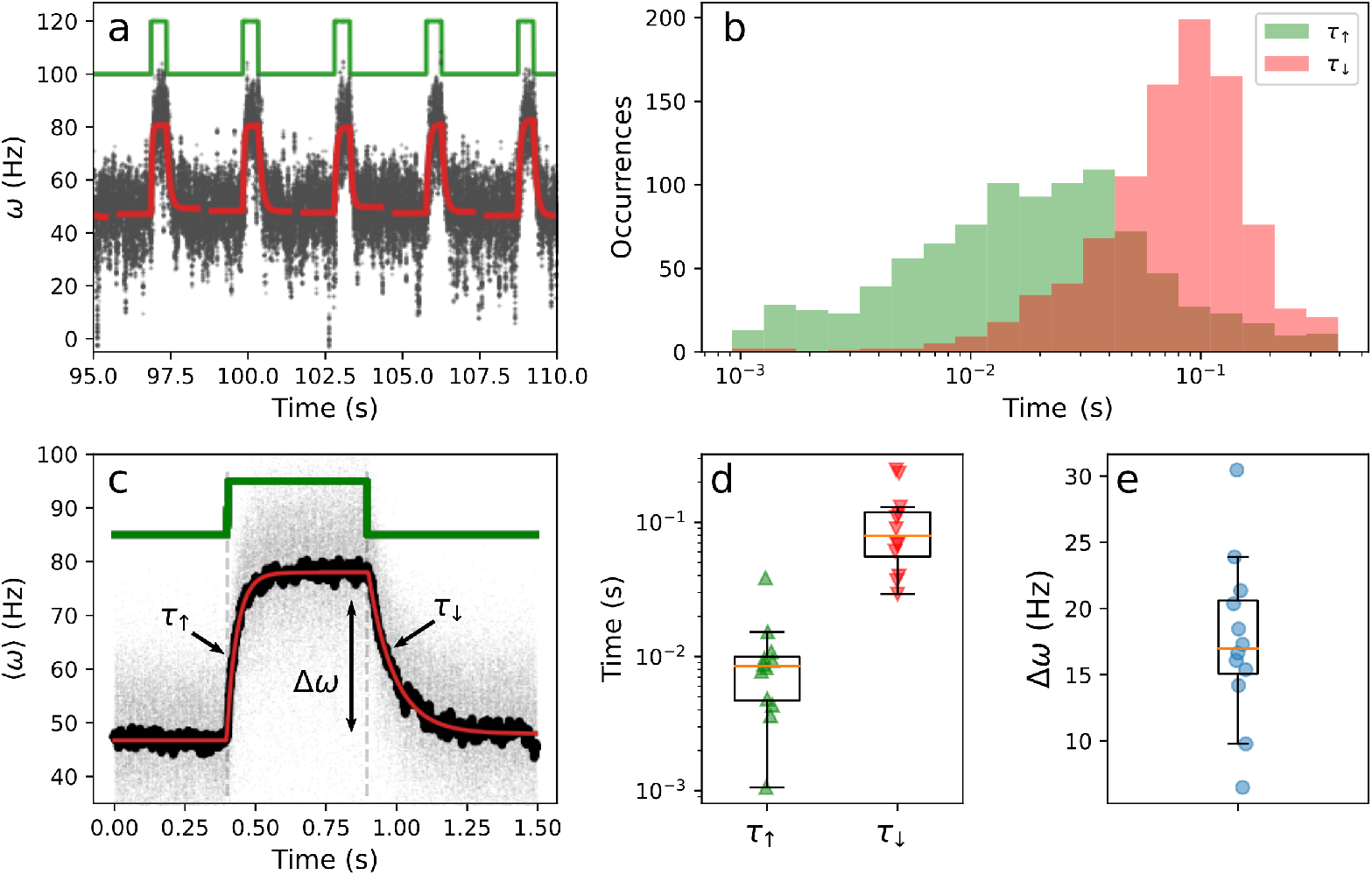
Global statistics. (a) BFM speed response of a *E. coli* expressing PR (grey) induced by a train of on-off rectangular laser pulses (green). A piecewise function composed of single exponential functions was fit to each speed transition generated by a laser pulse (red). Same as in Fig. 1e of the main text. (b) Histogram of the characteristic times, *τ*_↑_ (green) and *τ*_↓_ (red), extracted from 12 different motors and a total of 936 speed transitions. (c) Overlay of hundreds of laser-synchronised transitions (grey), mean of the transitions (black), and fit of the mean (red). Same as in Fig. 1e of the main text. (d) characteristic times *τ*_↑_ (green) and *τ*_↓_ (red) extracted from the average transition fit of 12 motors. (e) Increase in speed extracted from the average transition fit of 12 motors

### 5 Estimates of cellular electrical parameters from circuit analysis

Walter et al [1] estimate *V_r_* from the energy of NADH oxidation under physiological conditions (Δ*G* = 212 kJ/mol, with a stoichiometry of 6*H*^+^ per electron pair) as *V_r_* ≃ 360 mV. We further set *R_r_* = *βR_s_*, and use the circuit expressions 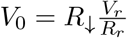 and 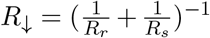 to obtain

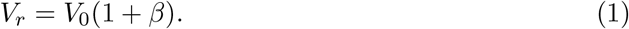

In the following, we assume i) a linear relationship between PMF and BFM speed (with zero y-intercept), and ii) that the PMF of energized cells is ~ 150 mV [2] prior to oxygen reduction and with a BFM rotating at ~ 200 Hz. Therefore, the experimentally measured steady state speed after one to two hours in a sealed flow cell, *ω*_0_ ≃ 50 Hz, and the speed under PR excitation *ω*_1_ ≃ 80 Hz (Fig. 1 of the main text), correspond to a voltage *V*_0_ = 37.5 mV and *V*_1_ = 60 mV, respectively. From Eq. 1, we then obtain *β* ~ 9. From *τ*_↑_ = *R*_↓_*C* we obtain

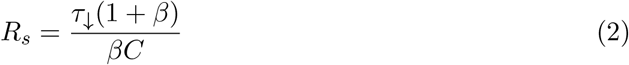

Using the measured value of *τ*_↑_ ≃ 100 ms, and the published value of *C* ≃ 10 ^14^*F* [1, 3], we get *R_s_* ≃ 1 × 10^13^ Ω, and *R_r_* ≃ 9 × 10^13^ Ω (we note that the assumptions made in [1] lead to *R_s_* ≃ *R_r_* ≃ 1 × 10^14^ Ω to 1 × 10^15^ Ω). Using the circuit expressions 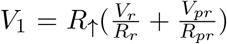 and 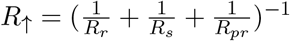, we obtain the expression for *V_pr_* as

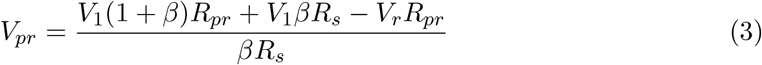

Using *R_pr_* ≃ 1 × 10^12^ Ω (obtained from the expression 1/*τ*_↑_ – 1/*τ*_↓_ = 1/(*R_pr_C*), see main text), we finally obtain *V_pr_* ≃ 62 mV.

### 6 Analytical solution to the cable equation

The 1D cable equation for the quantity *A*(*x, t*) (which can be voltage or concentration) can be written as function of space, *x*, and time, *t*, normalized to the characteristic length, *λ*, and characteristic time, *τ*, as,

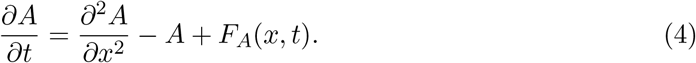

When the external source *F_A_* is applied at a point *x_o_* along a cable of length *L*, during the time window (*t*_1_ = 0, *t*_2_), one can write *F_A_*(*x, t*) = *δ*(*x* – *x_o_*)[*H*(*t*) – *H*(*t* – *t*_2_)], with *H*(*t*) the unit step function. The analytical solution (shown in Fig. 4 of the main text) can be written as [4]

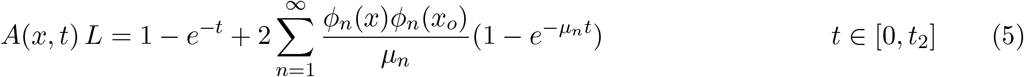

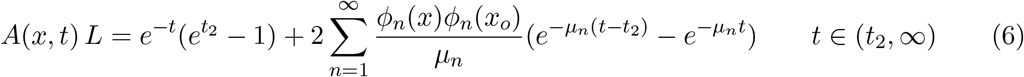

with *φn*(*x*) = cos(*nπx/L*) and *μ_n_* = 1 + *n*^2^*π*^2^/*L*^2^, (*n* = 1,2,..). Although the above solution was found in the context of cable theory [4], it can be applied to the diffusion problem, upon substitution of the relevant parameters.

### 7 Circuit and diffusion simulations

Here we show the results of the simulation of i) a simple circuit which models the spatial extension of the cell, and ii) of the diffusion equation with losses.

i) The circuit (SI Fig. 5a) is composed by two parallel *RC* sub-circuits, one of which includes the light-driven source. We consider both a voltage source *V_pr_* with internal resistance *R_pr_*, and a current source *I_pr_*. The two sub-circuits correspond to two spatially separated parts of the cell (where two sensors can be located), connected by the cytoplasmic resistance *R_i_* (internal). *R_s_* is the membrane resistance (sinks), *C*_0_ and *C*_1_ (of the same value) indicate the membrane capacitance at the two locations where the voltage measurement is performed. To compare the results obtained with the two sources (voltage and current), we fix *I_pr_* = *V_pr_*/(*R_pr_* + *R_eq_*), where *R_eq_* = 1/(1/*R_s_* + 1/(*R_s_* + *R_i_*)) is the equivalent resistance seen by the source. In SI Fig. 5b, the values of the electrical components are the ones obtained from the analysis described in the main text (see figure legend). We assume a cylindrical cell of 0.5 *μ*m radius, and a distance between the two sections of the cell, modeled by the two sub-circuits, of 20 *μ*m (in line with our measurements). In particular, this results in a cytoplasmic resistance *R_i_* = 10^7^ Ω, a value orders of magnitude smaller than the other resistances. With such small *R_i_*, SI Fig. 5b shows that the two sub-circuits respond identically, as the voltage across *C*_0_ and *C*_1_ (continuous and dashed lines of both colors, respectively) perfectly overlap. Using a voltage source (blue continuous and dashed lines) introduces an asymmetry in the characteristic times (*τ*_↑_ < *τ*_↓_, defined in the main text), which is not present with the current source (green continuous and dashed lines). To observe a difference between the transmembrane voltage measured across *C*_0_ and *C*_1_, we have to artificially increase *R_i_* by several orders of magnitude. This results in the traces shown in SI Fig. 5c, where the voltage across *C*_1_ (green and blue dashed lines, for both source types) reaches a lower plateau than the one across *C*_0_.

ii) In SI Fig. 5d,e we numerically integrate the 2D diffusion equation for the concentration *ρ*(*x, t*) with losses *k_s_* and sources *k_pr_*

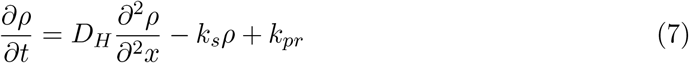

where *D_H_* = 9·10^-9^*m*^2^/*s* is the Grotthuss diffusion coefficient. The integration is performed on a cylinder (periodic boundary conditions on one direction and zero Neumann on the other) of 35 *μ*m length and 0.5 *μ*m radius (which effectively makes the system 1D), where the source k*pr* is spatially localized on one pole on the first 6 *μ*m of the cell, and is temporally switched on at *t*_0_ = 0 and off at *t*_1_ = 0.3 s. The concentration is monitored at two points on the membrane, indicated by circles labeled *C*_1_, *C*_2_ in SI Fig. 5d, the first at the location of the source, and the second at a distance of 20 *μ*m, to simulate the two farthest BFMs found in our measurements. The green continuous and dashed lines in SI Fig. 5e indicate the evolution of the concentration at *C*_0_ and *C*_1_, respectively, for a source modeled by a constant rate (*k_pr_*(*x, t*) = *const*. > 0, for *x* at the excited pole and *t* ∈ [*t*_0_, *t*_1_], otherwise *k_pr_*(*x, t*) = 0). We also introduce a Hill-Langmuir saturation mechanism in the source, to make *k_pr_* a decreasing function of concentration *ρ*, using

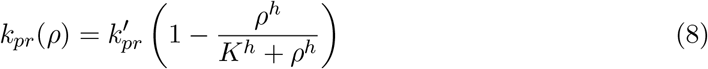

where 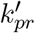 is the source rate at low concentration, *K* the equivalent of a dissociation constant, and *h* the Hill coefficient. In SI Fig. 5e, the blue continuous and dashed lines correspond

**Figure 5:**
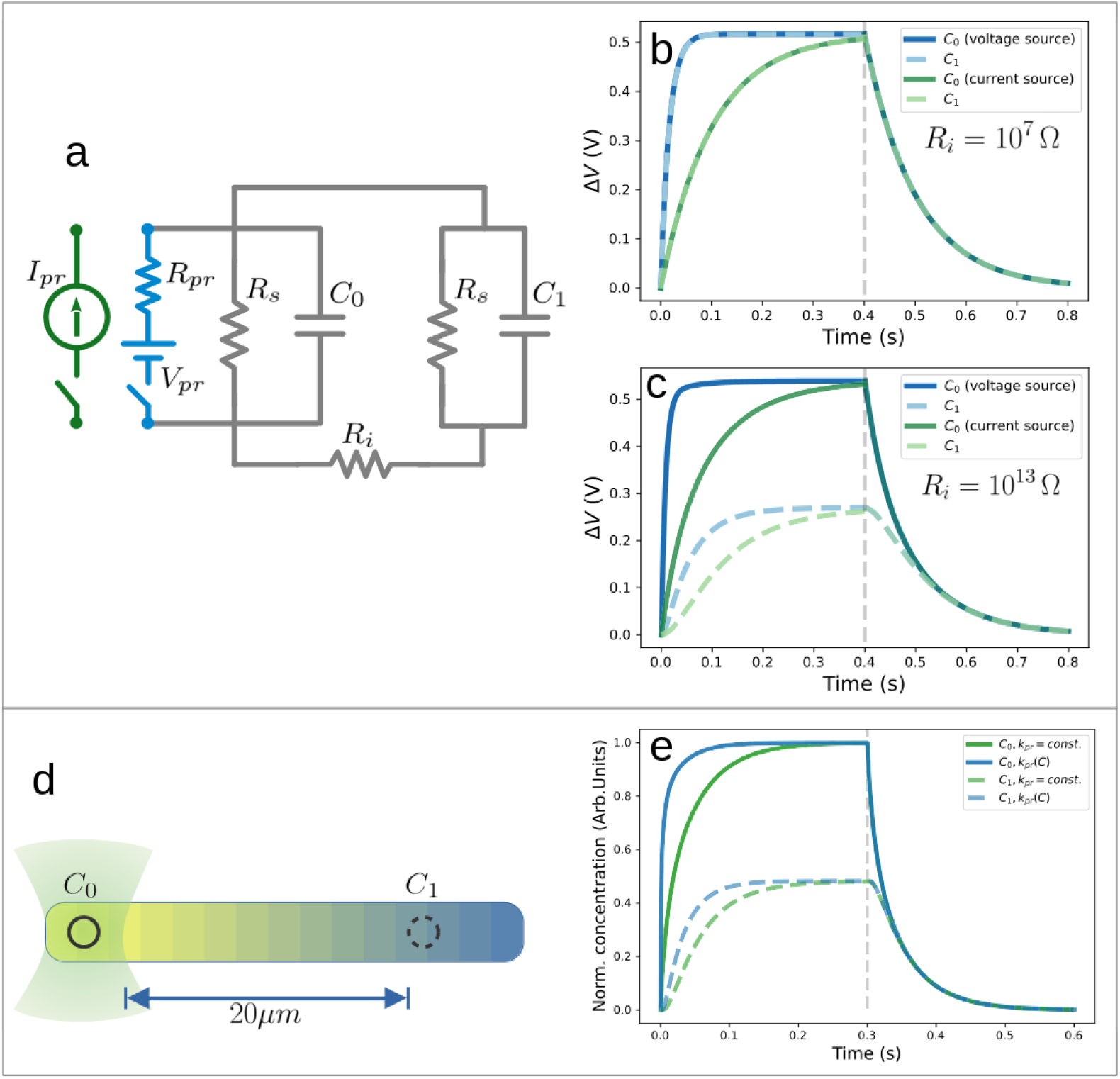
Circuit and diffusion simulations. (a) The circuit simulated is composed by two RC sub-circuits (membrane resistance from sinks *R_s_*, and membrane capacitance *C*) connected by an internal cytosol resistance *R_i_* (while the outer medium is considered conductive). We consider either a voltage (*V_pr_, R_pr_*, blue) or a current source (*I_pr_*, green). (b) Circuit simulation with parameters obtained from the analysis described in the main text, where we fix *R_i_* = 10^7^ Ω (the resistance of a cylindrical cell of radius 0.5μm, length of 20 *μm*, and volume resistivity of 100 Ω cm [3, 5]). The parameters used are *R_s_* = 10^13^ Ω, (so *R_i_* ≪ *R_s_*), *C*_0_ = *C*_1_ = 10^-14^ *F*. The blue lines (continuous and dashed) correspond to the voltage source with *V_pr_* = 0.62 V, and *R_pr_* = 10^12^ Ω. The green lines (continuous and dashed) correspond to the current source, with *I_pr_* = *V_pr_*/(*R_pr_* + *R_eq_*), where *R_eq_* = 1/(1/*R_s_* + 1/(*R_s_* + *R_i_*)) is the equivalent resistance of the gray circuit in a). The continuous lines (blue and green) correspond to the voltage across *C*_0_. The dashed lines (blue and green) correspond to the voltage across *C*_1_. The voltage source introduces an asymmetry in the characteristic times (*τ*_↑_ < *τ*_↓_), which is absent with the current source (*τ*_↑_ = *τ*_↓_). (c) Same as (b), but with an increased internal resistance (*R_i_* = 10^13^ Ω, so *R_i_* = *R_s_*), which produces a reduction of the value of the plateaus reached by the voltage across *C*_1_ (dashed lines), for both source types. (d) Diffusion simulations. Eq. 7 is integrated on a cylinder with diffusion coefficient *D_H_* = 9 · 10^-9^ m^2^/s and *ks*=20 s^-1^. The source is located at one pole of the cell, and it is switched on for 0.3 s. The concentration is monitored at the two points *C*_0_, *C*_1_. (e) Evolution of the normalized concentration. Green lines correspond to the choice *k_pr_* = *const*. = 1 (see text), while the blue lines correspond to the choice *k_pr_* = *k_pr_*(*ρ*) (Eq. 8, with 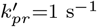, *K* = 10^-4^ and *h* = 2). Continuous and dashed lines indicate the measurement at point *C*_0_ and *C*_1_, respectively. An asymmetry in the characteristic times (*τ*_↑_ < *τ*_↓_) appears with the choice *k_pr_* = *k_pr_*(*ρ*) (blue lines). A clear difference is visible between the plateaus reached by *ρ* at *C*_0_ and *C*_1_. A difference in the plateau reached at *C*_1_ with respect to *C*_0_ is present, as for the electric circuit with low internal resistance in (c).

to such choice of *k_pr_*(*ρ*). As in the case of the voltage source in the circuit above, this mechanism creates an asymmetry in the characteristic times (*τ*_↑_ < *τ*_↓_) which is absent in the case of constant source rate.

Among the scenarios discussed above, our measurements of the spatial dynamics of the PMF are only compatible with SI Fig. 5b with a voltage source, where the voltage measured at the two distant sensors evolves identically, with asymmetric characteristic times. This is achieved by the voltage in the presence of a low cytoplasmic resistance *R_i_* (with respect to the other resistances of the circuit) in the circuit model, and cannot be reproduced either by a circuit with *R_i_* of the same order of the other resistances, or by diffusion of concentration *ρ*. In the case of diffusion, even considering the highest diffusion coefficient and reduced dimensionality (2D or 1D), the concentration develops a spatial gradient along the cell which produces, once measured by two sensors placed at a distance of 20 *μ*m, traces with distinctly different plateaus, in contrast with our observations.

## Notes

### Competing Interest Statement

The authors have declared no competing interest.

